# Event segmentation reveals working memory forgetting rate

**DOI:** 10.1101/571380

**Authors:** Anna Jafarpour, Elizabeth A Buffalo, Robert T Knight, Anne GE Collins

## Abstract

We encounter the world as a continuous flow and effortlessly segment sequences of events into episodes. This process of event segmentation engages working memory (WM) for tracking the flow of events and impacts subsequent memory accuracy. WM is limited in how much information is retained (i.e., WM capacity) and for how long the information is retained (i.e., forgetting rate). It is unclear which aspect of WM limitations affects event segmentation. In two separate experiments with multiple tasks, we estimated participants’ WM capacity and forgetting rate in a dynamic context and evaluated their relationship to event segmentation. The results across tasks show that individuals who reported more movie segments than others (fine-segmenters) have a faster decaying WM. A separate task assessing long-term memory retrieval reveals that the coarse-segmenters have better recognition of temporal order of events in contrast to the fine-segmenters who performed better at free recall. The findings show that event segmentation employs dissociable memory strategies and depends on how long information is retained in WM.

## Introduction

Event segmentation is the process of discretizing a flow of events into episodes (Zacks et al., 2007; Zacks and Swallow, 2007). It is done naturally even without instructions (Jafarpour et al., 2019). Event segmentation is thought to be fundamental in shaping cognitive processes such as working memory (Kurby and Zacks, 2008b; Radvansky, 2017; Richmond et al., 2017). Current event segmentation models posit that working memory tracks the flow of events to determine the segments and integrates information and retains ‘what is happening now’ (called event models) within an event segment (Kurby and Zacks, 2008a; Sargent et al., 2013). An event segment is thought to end when a perceived event does not match the expectations of what would proceed from a prior flow of information (Bein et al., 2020; Franklin et al., 2020). However, working memory limitation may contribute to event segmentation. A working memory system that fails to retain information must update the event model often or utilize other strategies to compensate for its limitations. Subjects who segment the flow of an event as defined by the experimenter have a better working memory performance than those with a discrepancy in event segmentation (Karuza et al., 2019; Sargent et al., 2013).

It is not clear what aspect of working memory is linked to event segmentation. We study the link between the number of determined events and working memory limitations. We focus on two independent aspects of working memory indexing the amount of information retained and the duration of maintenance (Baddeley, 2012, 2003; Bays and Husain, 2008; D’Esposito and Postle, 2015; Oberauer et al., 2018). The first factor, how much information can be held in memory, defines working memory capacity (Bays et al., 2009; Burgess and Hitch, 1992; Cowan, 2010; Vogel and Machizawa, 2004). The other factor, how long the information persists in the face of time and interference, is measured as the working memory forgetting rate (Baddeley, 2012; Collins and Frank, 2012). A challenge in measuring these limitations is that other memory systems, such as long-term memory, can also contribute to the accuracy of working memory (Jafarpour et al., 2017; Rose et al., 2016; Zokaei et al., 2014). In this study, we implemented a working memory/association learning task, designed by Collins and Frank (2012), that disentangles the working memory and learning system including episodic memory. The task requires learning associations between images and actions in blocks of variable image set sizes (Figure 1B and 1C). This task allowed us to computationally estimate the working memory capacity and forgetting rate in each participant. We modeled the association learning behavior with a dynamic mixture of learning and working memory components (adapted from Collins et al., (2017)). This reinforcement learning and working memory (RLWM) model and the task characterize working memory used in dynamic contexts that involve multiple memory systems. For that reason, we hypothesized that it would be suitable for relating to potential working memory use in dynamic naturalistic experiments where multiple memory systems may also be engaged, such as naturalistic event segmentation.

**Figure 1.**
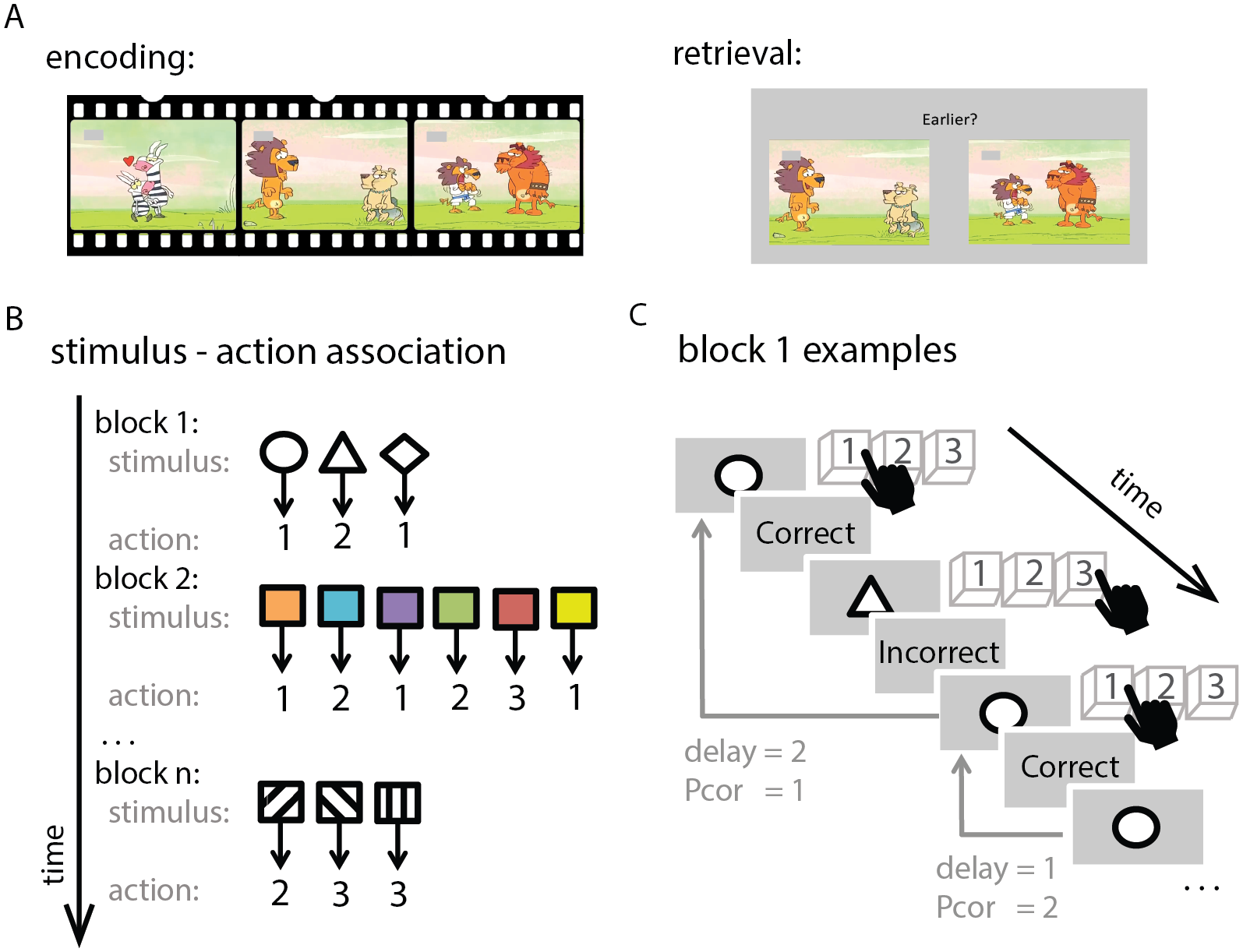
Experimental design: The experiment consisted of four parts, the first two of which are depicted here. (A) Temporal memory test: At encoding (left), participants watched three muted movies (frames from one movie is shown). At retrieval (right), participants saw two movie frames and determined their temporal order by pressing the left or right key. There were 35 temporal order questions per movie. (B) Association learning task: Participants performed a block-design association learning task. In each block, participants learned the association between a set of images and actions by trial and error. There were three possible actions (key 1, 2, or 3) and feedback was provided. The image set size varied across blocks, ranging from 2 to 6. (C) Three example trials. ‘Delay’ parameter quantifies the number of intervening trials from the last time the stimulus was encountered, and ‘Pcor’ quantifies the number of trials that the action choice was correct. The last two parts of the experiment were segmentation and free recall tasks.

The RLWM model parameterizes working memory limitations as the probability that the stimulus-action association can be kept in working memory and can be used accurately for choosing the rewarding associated action in the future. The association learning task manipulates working memory limitation with discrete set sizes and delays; hence, the model approximates the working memory resource limitation to the number of retained associations, akin to the slot model, ranging the capacity from 2 to 6 (Cowan, 2000; Ma et al., 2014). Accordingly, the probability that the correct stimulus-action association is available in working memory depend on the working memory capacity C and set size ns. Specifically, if the set size is within capacity, the model makes maximal use of information in WM, but if the set size is larger, information in WM contributed less to choices, according to the probability 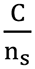, that items were stored in WM. The working memory forgetting rate in RLWM can also approximate how fast the memory of the retained association is forgotten when facing new stimuli. The forgetting rate indicates how fast the probability of selecting a correct association decayed to the chance level (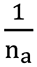, where n_a_ = 3 is the number of possible actions).

Traditionally, event segmentation has been studied with experimentally defined boundaries, where a change in an associated stimulus, such as the background image or the semantic category of a sequence of images determines a new segment (DuBrow and Davachi, 2013; Ezzyat and Davachi, 2014a), or the expected event boundaries are defined by the experimenter (Sargent et al., 2013; Speer et al., 2007, 2003). In these cases, event segmentation is discussed as the agreeability of an individual’s segmentation with the predefined event boundaries. The empirical data, however, show that the segmentation of a naturalistic sequence of events is more subjective. Although event segmentation can be modulated to a fine or coarse grain through instructions (Sargent et al., 2013; Speer et al., 2007), across a population, individuals differ in how many events they identify (Jafarpour et al., 2019; Sargent et al., 2013; Zacks et al., 2006). For instance, identified event boundaries for a naturalistic flow of events varies across subjects viewing the same stimuli (the maximum agreeability in Speer et al. (2003) was at 80% (n = 11) and in Jafarpour et al. (2019) was at 60% (n = 80)). Here, we investigated the source of individual variability in event segmentation without assuming predefined event boundaries.

We hypothesized that sub-components of memory are linked to the number of identified boundaries in a flow of events (i.e., the number of events). One possibility is that infrequent segmentation overwhelms working memory storage, resulting in faster forgetting and poor subsequent memory of events. Another possibility is that limited working memory constrains information retention leading to frequent segmentation and utilization of other memory systems such as long-term memory. It is evident that episodic memory (DuBrow and Davachi, 2013, 2014; Ezzyat and Davachi, 2011, 2014b; Tubridy and Davachi, 2011) and utilizing stories (a.k.a. scripts or event schemata) (Keidel et al., 2017; Zwaan and Radvansky, 1998) are involved in event segmentation. Event boundaries are more memorable than other events and remembering the temporal order of events across event boundaries is more difficult than within the boundaries (Heusser et al., 2018; Horner et al., 2016). We predicted that frequent event segmentation would enhance long-term memory and diminish temporal order memory.

We studied the link between the number of determined events and working memory limitations by administering two independent tasks (a movie segmentation task and a working memory/association learning task) to a group of healthy participants (Figure 1). Participants watched a movie and were subsequently tested on their memory. They later watched the same movie and reported subjective event boundaries. We allowed the individual to utilize their natural strategy for event segmentation. All movies were novel (sound off) animations with simple illustrations (Figure 1). We selected movies with different storylines. The storyline of one of the movies was non-linear so that interchanging the epochs of the movie did not affect the story, another movie had a linear storyline with non-interchangeable epochs, and the third movie had a mixed storyline (see Supplemental Information for the storylines). Including both linear and non-linear storylines allowed us to observe whether individual variability in event segmentation is due to utilizing stories, such as schematic knowledge of the story (Bower et al., 1979), or whether segmentation variability is independent of story knowledge (Sargent et al., 2013). The study included the association learning task with variable association set sizes (Collins et al., 2017). We used the described RLWM model to estimate the participant’s WM capacity and forgetting rate. Participants also performed a temporal order recognition task and wrote a paragraph about the movies, i.e., a free recall task. We ran this study twice: experiment 1 was performed in the lab with a controlled experimental setting. Experiment 2 was a replication of experiment 1 in an online setup.

## Results

Behavioral results for individual tasks were consistent with previous studies (see supplementary info). In particular, we replicated the established observation that RLWM provided the best fit compared to reinforcement learning (RL) models without a working memory component (Figure S2). We focus here on cross-task results.

In experiment 1, the cross-task comparison showed a U-shaped relationship between the variation in the total number of determined events in the movies and the working memory forgetting rate (p < 0.001; Figure 2; Table S3), but there was no linear or quadratic relationship between working memory capacity and the number of events (mean capacity = 3.38 (SD = 1.10); linear fit p = 0.64; quadratic fit p = 0.89). Across the participants, the total number of determined events was between 2 and 148 events (M = 38, SD = 27.80). Excluding the participant with the outstanding number of events (148) did not change the results (Table S2; Figure S1 for the agreeability of event boundaries for the movies; also see the supplemental information for additional details). The U-shaped relationship between forgetting rate and number of events was observed in all three movies with different storylines (p < 0.001; Figure S4). Adding the quadratic term explained the data better than a linear fit, accounting for additional complexity via Akaike Information Criterion (AIC; Table S1-S3). Three participants were excluded due to low fit quality (see Supplemental Results); however, including these participants did not change the main results (Table S1-S3).

**Figure 2.**
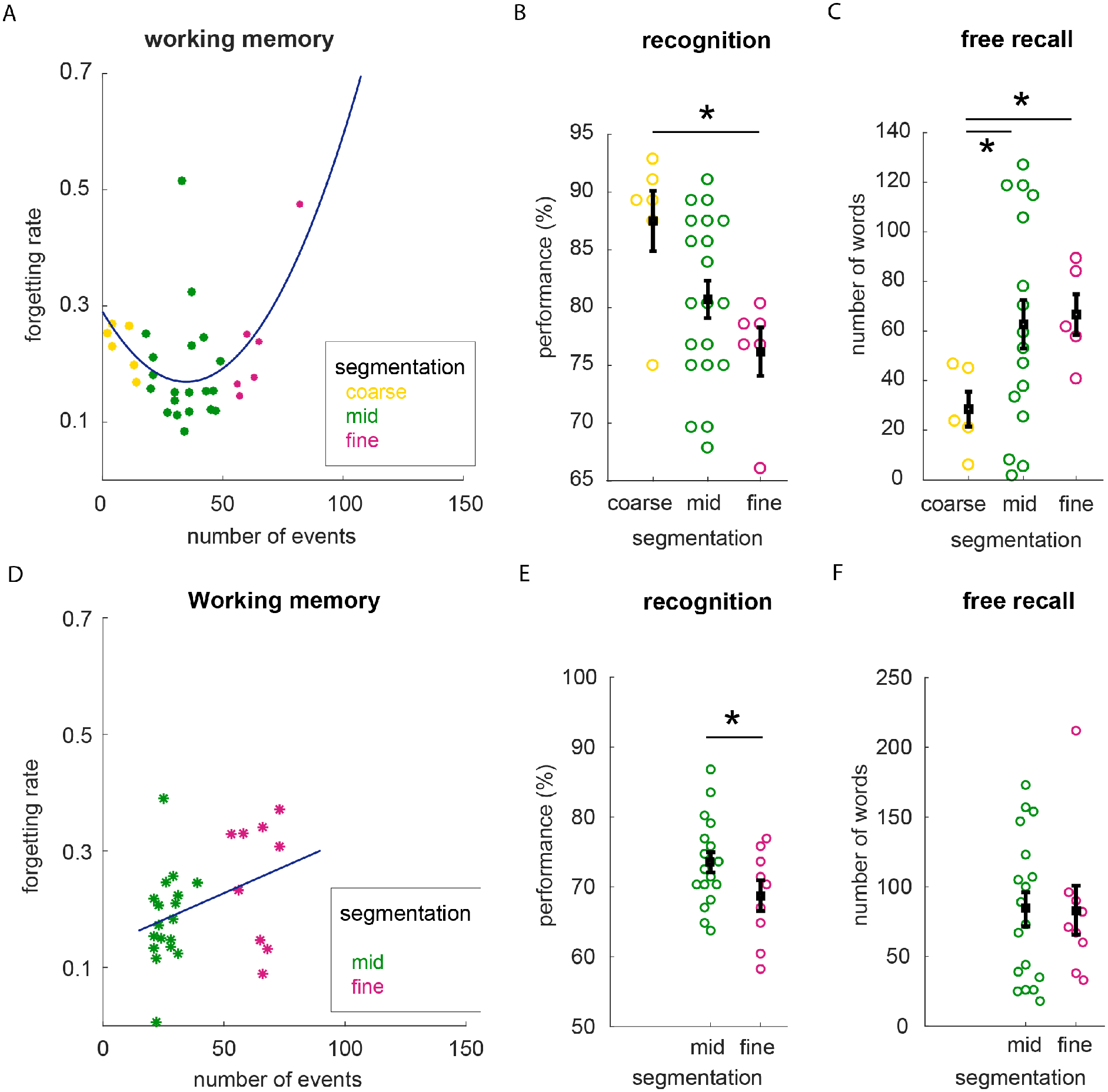
Cross-task results (A-C) Experiment 1: (A) The number of determined events and working memory forgetting rate had a U-shaped relationship. The quadratic regression is fit without the outlier. (B) The coarse-segmenters showed better recognition of the temporal order of events than fine-segmenters. (C) In contrast to order recognition, the fine-segmenters recalled more events about the movies than the coarse-segmenters. (D-F) Experiment 2: (D) The working memory forgetting rate correlated with the number of determined events. (E) The coarse-segmenters showed better recognition of the temporal order than the mid group. (F) There was no significant difference between mid and fine segmenters in the number of recalled words. Error bars represent the standard error of the mean. Each dot depicts a participant. (* p < 0.05).

The U-shaped relationship suggests that individuals with a faster working memory forgetting rate used at least two different strategies for segmentation (fine and coarse) and may differentially utilize long-term memory. We used the total number of determined events to group the participants into fine -, mid-, or coarse-segmenters, depending on whether the number of reported events was one standard deviation more than the mean number of events (fine), one standard deviation less than the mean number of events (coarse), or within one standard deviation of the mean (mid). This grouping allowed us to study the cross-task results in more detail. Follow-up tests compared the behavior of fine- and coarse-segmenters to understand the underlying differences between their behaviors.

We observed that the recognition and recall performance of the participants who segmented too many or too few events were different. The fine-segmenters had a longer response during the free recall task (M = 66.66; SD = 19.97 number of words) than the coarse-segmenters (M = 28.46; SD = 17.23; two-sample t-test t(8) = −3.237, p = 0.011; Figure 2C). There was a trend towards a correlation between the number of events and the number of words at free recall for all the movies (Spearman r = 0.339, p = 0.089; note that there was no presumed number of events in the movies to subsequently count the number of recalled events.) We did not observe any false memories or intrusions. We further assessed the free recall performance with a natural language processing algorithm that determines the number of “realis” words used for factual events (Sap et al., 2020; Sims et al., 2019). Although this method does not include the recalled details of the movies, we observed a trend toward greater recall of factual events by fine-segmenters than coarse segmenters (two-sample t(8) = −1.69, p = 0.06; see the supplemental results for the main effect of movie type and the number of recalled words).

In contrast to fine-segmenters, the coarse-segmenters performed better at the temporal order recognition for the linear movie (coarse-segmenters: M = 87.5, SD = 2.6; fine-segmenters: M = 76.1, SD = 2; two-sample t(10) = 3.32, p = 0.004; Figure 2B; for all movies: t(10) = 1.97, p = 0.03), although the inference about the temporal order of the movies was accessible to both groups (Lositsky et al., 2016). Across all participants, temporal order recognition accuracy tended to decrease with an increased number of determined events (linear movie: r = −0.34, p = 0.053; all movies: r = −0.25, p = 0.15). Accordingly, we propose that the coarse-segmenters utilized the overall story of the movies to compensate for fast forgetting. This strategy reduced memory details resulting in coarse-segmenters having a worse free recall performance than the fine-segmenters (See the supplemental results for the overall temporal order memory and free recall performances; there was no significant correlation between free recall and temporal recognition memory performance; r = −0.19, p = 0.35.)

In experiment 2, we sought to replicate the link between working memory forgetting rate and event segmentation in an independent group of people who performed the study online due to Covid-19 pandemic restrictions. The online data collection was not conducted in a controlled environment. Therefore, we excluded participants who performed poorly in any parts of the experiment (see the supplementary results for details) to ensure analysis of participants who were fully engaged. In the online data set, the distribution of the number of events was narrower than in experiment 1. Specifically, no participants in experiment 2 fell in the range of course segmenters as defined in experiment 1. Participants in experiment 2 showed two distinct clusters in the number of determined events (Figure 2D and 2SB) corresponding to fine- and mid-segmenters. The fine- and mid-segmenters performed equally well in the learning task (for example, of the last trials in the easiest blocks; set size 2; t(25) = 0.17, p = 0.86).

Consistent with experiment 1, we observed a correlation between the number of determined events and working memory forgetting rate: the working memory forgetting rate was higher for people who reported a high number of events (r = 0.37, p = 0.05). These findings replicated findings from experiment 1 showing that the fine-segmenters had a faster working memory forgetting rate than mid-segmenters (one tailed t-test: t(25) = −1.9, p = 0.03). The mid-segmenters recalled the order of events with a higher accuracy than the fine-segmenters (t(25) = 1.84, p = 0.03). There was a trend towards a correlation between the number of determined events and temporal order memory (r = −0.35, p = 0.06). We did not observe any difference in recall length or the number of factual words (i.e., the realis) between mid- and fine-segmenters, consistent with findings from experiment 1. Generally, the recall length was longer in experiment 2 (where coarse-segementers were not found) than in experiment 1 (t(54) = 4.18, p < 0.001; Figure 2C and 2F) and more realis events were recalled in experiment 2 than in experiment 1 (t(54) = 4.5, p < 0.001).

## Discussion

In two independent studies, we observed a link between event segmentation, working memory forgetting rate, and subsequent memory performance. We used a newly-established task measuring the use of working memory in a dynamic decision-making context (Collins and Frank, 2012) to estimate the participants’ working memory capacity and forgetting rate. Using cross-task comparisons, we found that fine- and coarse-segmenters have a faster working memory forgetting rate than the mid-segmenters and they employ long-term memory strategies to compensate for the fast working memory decay.

Previously, Sargent et al. (2013) showed that event segmentation correlates with working memory performance in reading span, operation span, and word-list memory tasks (Daneman and Carpenter, 1980; Turner and Engle, 1989). In that study, event segmentation was assessed as the discrepancy from an experimentally assumed event boundary. In that case, the fine and coarse segmentation would have been characterized with a low segmentation ability and poor working memory. The adopted working memory tests in the previous study, however, could not clarify what aspect of working memory limitation is linked to event segmentation. In this study, we found a link between event segmentation and working memory forgetting rate. The link suggests that event segmentation is related to the retention of the flow of events, rather than the retention of information which would be limited by working memory capacity.

At encoding, perception of event boundaries naturally engages the memory network (Jafarpour et al., 2019) and increases recall performance (Newtson and Engquist, 1976; Pettijohn et al., 2016; Sols et al., 2017). Furthermore, people remember the event-boundaries better than other events (Newtson and Engquist, 1976). It is previously shown that a fine-grained segmentation strategy benefits source memory (Hanson and Hirst, 1991, 1989; Heusser et al., 2018). However, with fine segmentation, the contextual representation of a sequence is updated resulting in less access to the temporal memory of events that occurred across event boundaries, e.g., “walking through doorways” (DuBrow and Davachi, 2014; Pettijohn and Radvansky, 2016). Congruent with this previous work, we observed that the fine-segmenters have worse performance in the temporal order recognition task than coarse- or mid-segmenters who have a better recall performance.

Computational models of event segmentation suggest that sequences are segmented based on predictions from prior knowledge about sequences of events, perhaps by utilizing stories (Hsieh et al., 2014; Schapiro et al., 2013; Zacks et al., 2011), or the sequences of events are linked to a temporal context that changes with event boundaries (DuBrow and Davachi, 2014; Franklin et al., 2020; Howard et al., 2005; Lositsky et al., 2016; Radvansky and Zacks, 2011; Zacks and Swallow, 2007). Indeed, a change in the story or temporal context with segmentation leads to less accessibility of memory of temporal order of events across boundaries, and retaining the temporal context benefits temporal memory (DuBrow and Davachi, 2014; Horner et al., 2016; Manning et al., 2011). Here we showed that working memory forgetting rate also plays a role in event segmentation. In a naturalistic setting, the retention can be fulfilled by utilizing stories or relying on a working memory system with a slow forgetting rate.

An outstanding question concerns the causality of the relationship between working memory and event segmentation. One possibility is that working memory is a primary cognitive mechanism and event segmentation is determined by limits in working memory. Accordingly, participants with a faster forgetting rate use alternative mechanisms such as utilizing scripts or long-term memory to compensate. An alternative possibility is that event segmentation is a primary cognitive mechanism (Ongchoco and Scholl, 2019; Radvansky, 2017; Richmond et al., 2017). In this case, participants who segment more often lose access to the information from the previous events (Ezzyat and Davachi, 2014b; Horner et al., 2016) leading to a fast forgetting rate or to relying on storylines to keep a track of what happened. A third possibility is that both working memory and event segmentation engage common cognitive mechanisms, such as utilizing stories or scripts. Utilizing a script facilitates memory (Farag et al., 2010; Gobet et al., 2015; Sargent et al., 2013; Zacks et al., 2010). For example, a phone number’s schema enables effective segmentation and memory for a 10-digit number (Miller, 1956).

In conclusion, we observed that the working memory forgetting rate reflects individual differences in event segmentation. A relationship between the number of determined events on one task and the forgetting rate on another task suggested that participants with a faster forgetting rate used two different strategies during encoding and retrieval: utilizing scripts versus relying on episodic memory. Taken together, these data suggest that working memory plays a key role in shaping event segmentation, and people with faster working memory forgetting rates utilize alternative cognitive processes for encoding and retrieval of events.

## Limitations of the Study

A limitation of this study is that we did not detect the coarse-segmenters in experiment 2 when the data was acquired online to mitigate the COVID-19 pandemic. Coarse-segmentation may be a strategy that was not adopted because of the change in the experimental setup (in the lab versus online) and the absence of the experimenter supervision during data acquisition. Another possible reason for the lack of coarse-segmentation in the online data is that a smaller percentage of the population are coarse-segmenters and by chance, they were not sampled in the online data set. Future lab experiments on individual differences in event segmentation can clarify why this was the case.

The observed individual variability cannot be explained by the participants’ level of task engagement. Individuals who identified fewer events – potentially a less motivated group (coarse-segmenters) – performed better in the memory recognition test. In addition, all participants’ memory and learning performances were above chance. Nevertheless, additional studies will be needed to determine the scope of the difference between fine- and coarse-segmenters in utilizing stories and free recall. For example, is the observed difference specific to long-term memory or does our individual difference result generalize to other behavior such as generating stories and imagination?

## Resource Availability

The data used to support the findings of this study are available from the corresponding author upon request.

## Acknowledgments

We thank Cassandra Lei, Crystal H. Shi, and Megan Schneider for experiment 1 data collection and Amy Zou for experiment 2 data collection. We thank Maarten Sap for realis annotations. This research was supported by the National Institute of Health, National Institute of Mental Health, K99MH120048-01 (A.J.), and the National Institute of Neurological Disorders and Stroke, 1U19NS107609 (E.A.B.) and R37NS21135 (R.T.K.).

## Author Contributions

Conceptualization Ideas: A.J.; Methodology: A.J.; Software: A.J. and A.G.E.C.; Validation: A.J. and A.G.E.C.; Formal Analysis: A.J. and A.G.E.C.; Investigation: A.J.; Writing – Original Draft: A.J.; Writing – Review & Editing: A.J., E.A.B., R.T.K., and A.G.E.C; Visualization: A.J.; Supervision: E.A.B., R.T.K., and A.G.E.C; Funding Acquisition: A.J., E.A.B., and R.T.K.

## Declaration of Interests

The authors declare no competing interests.

## 1 Supplementary material

## 2 Participants

Experiment 1: 36 healthy and English-speaking adults (25 female) were recruited through the online University of Berkeley Psychology Department Research Participation Program. Participants provided informed consent and were compensated ($36 or 3 course-credits). The Office for the Protection of Human Subjects of the University of California, Berkeley approved the study protocol. The mean age was 20.3 (SD = 1.9) and ranged from 18 to 27. 29 of 36 participants performed the additional free-recall task (the data from the first 7 participants was not recorded due to a technical issue). All participants were right-handed by self-report. We discarded one participant because she identified two standard deviations more events than the group average but including her did not change the results. The minimum temporal order recognition accuracy for the linear movie was at 66% and for the non-linear movie was at 54%.

RLWM did not accurately model the performance of three participants. The estimated learning rate was too high (*α* > 0.9, two standard deviations larger than the mean; M = 0.16, SD = 0.33) and the estimated mixture weight was too low (*w*_0_< 0.6; M = 0.83, SD = 0.18) in the three participants, indicating that the working memory module of the model was not functioning in a regime representative of cognitive working memory function. AIC for one of these subjects also showed that RL4 was a better fit than RLWM for her. The three subjects were excluded from the analysis with forgetting rate estimations, although including them did not change the overall results (Table S1-S3).

Experiment 2: 101 healthy and English-speaking adults were recruited through the online University of Berkeley Psychology Department Research Participation Program. They provided an informed consent and received course credits for the participation. The study was run remotely by providing a link to the online task. Participant’s ran the study at their desired time of the day. 98% of participants performed above 70% for the last trials of learning trials for set size 2. 50% of participants wrote something about the movies. 65.3% of participants performed the order of events in the linear (easier) movie for at least as well as the worst participant performance in experiment 1 (at 66%). 19 participants did not indicate any events in the movie and two participants were excluded for outlier number of events. Accordingly, 76% of participants determined the number of events in a normal range. Cross-tasks, 27 of participants fulfilled all tasks. There were 19 female (8 male), age ranged from 18 to 22 (two participants did not report their age, mean = 20 years, SD = 1.2).

## 3 Experimental design

Experiment 1: The experiment ran on a desktop PC and a standard TFT monitor, in a sound-attenuated recording room. It consisted of four parts. First, participants watched three novel mute animations (each ∼3 minutes long) with differing storylines – linear (interchanging the movie epochs hurts the story), non-linear (interchanging the movie epochs did not hurt the story, like Tom and Jerry animation), and mixed (see the supplementary material for the stories). They were instructed to watch the movies carefully because we would ask questions about them. The types of storylines were identified by asking a group of naïve observers about the interchangeability of epochs of the movies.

There were 35 recognition memory tests per movie. At each test, subjects saw two scenes from a movie - located on the left and right sides of the screen for 2 seconds. Then a prompt appeared asking the participant to indicate the order of the scenes. Half of the prompts were asked which scene was ‘earlier’ or others asked which scene was ‘later’, in a pseudo-random trial order (Figure 1). Participants used left or right arrow keys to respond. The tested movie scenes were between 1.6 to 60 seconds apart for the mixed movie (the mean scene distance was 18.84 seconds, SD = 18.31), 1.6 to 62 seconds for the non-linear movie (M = 16.66, SD = 17.27), and 1.6 to 48 seconds for the linear movie (M = 15.02, SD = 13.22).

The participant then performed a version of an association learning task (Figure 1B and 1C) to evaluate working memory characteristics, namely working memory capacity and forgetting rate (Collins et al., 2017). In this task, participants used their dominant hand to select from three possible actions when they saw an image. The possible actions were pressing the J, K, and L keys from a keyboard (depicted as action 1, 2, and 3 in Figure 1). They used trial and error to learn the correct image-action association. The probability of an action being paired with an image was equal (1/3); thus, an action could pair with more than one stimulus. Participants learned the associations in 12 repetitions of each stimulus; the repetitions were pseudo-randomly interleaved. This procedure repeated in a block-design and included 22 blocks (3 blocks of set sizes 6, 5, and 4; 4 blocks of set size 3, and 6 blocks of set size 2). The stimulus set-size varied in each block to manipulate the requirement for capacity-limited and delay-sensitive working memory (Figure 1C). Participants studied the whole stimuli set at the beginning of each block.

The reward value for a correct response differed across stimuli; an incorrect response yielded no reward. Only one action for a stimulus was correct and each correct stimulus– action association was assigned a probability (p) of yielding a 2-point versus a 1-point reward, and this probability was either high (p = .80), medium (p = .50), or low (p = .20). We counter-balanced the probability within-participant and blocks to ensure an equal overall value of different set sizes and actions. Participants had 1.4 s to respond. The feedback was displayed for 0.5 s. There was an inter-stimulus interval of 0.5 - 0.8 s in which a fixation cross was shown.

After performing the association learning task, participants watched the movies again; this time they segmented the movies by pressing a spacebar to indicate the start of a new event. We instructed them to press a key “whenever something new happened.” We told the participants: “we want to segment this movie into episodes”, and we explained that segmentation could occur as often as they liked. The consistency of frequency of event segmentation for each participant was determined across the three movies. Finally, participants performed a surprise free recall test, where they were asked to write a paragraph describing the story of each movie. Participants were instructed not to worry about grammar and wording - simply “write what came to their mind.” The free recall test allowed us to investigate the relationship between the individual difference in event segmentation and subsequent memory performance.

Experiment 2: We replicated a subset of the experiment 1 online. Participants remotely ran the study on their personal computers. The study was written in javascript using jsPsych (de Leeuw, 2015). It included the association learning task, the encoding of the linear and non-linear movies, the recognition of temporal order of the movies, followed by the event segmentation task and free recall.

## 4 Analysis details

### 4.1 Association learning

We analyzed the association learning task in two ways, without modeling and with reinforcement learning and working memory (RLWM) modeling, consistent with previously published studies (Collins, 2018; Collins et al., 2017, 2014; Collins and Frank, 2018). Trials with missed responses or with less than 200 ms response time were discarded. To generate learning curves, we analyzed the proportion of correct choices as a function of the number of iterations (how many times the stimulus was encountered) and set size. Next, we used a multinomial logistic regression to evaluate the performance with respect to three parameters - the set size (number of stimuli in a block), delay (number of trials since the last correct choice for the current stimulus), and previously correct answers (number of correct choices made so far for the current stimulus) - and their interactions. We quantified the effect of working memory and learning on a trial-by-trial basis by modeling the probability of a correct choice for each participant as a function of the three parameters: set size, the number of previously correct answers, and delay (see Collins et al., (2017) for details).

### 4.2 Movie segmentation

We evaluated the keypresses during the movies and discarded any keypress that occurred less than 100 ms from the previous keypress to remove multi-clicks. Then we quantified the number of events by counting key presses for each movie. A ranked (Spearman) correlation was utilized to identify the consistency of individual differences across the movies.

### 4.3 Modeling

We fit three models to the trial-by-trial responses for each subject: two-parameters reinforcement learning (RL2), four-parameters reinforcement learning (RL4), and a modified RLWM model (Collins et al., 2017). We used the Akaike Information Criterion (AIC) to select the best model considering the number of parameters used in each model. RLWM was used as a baseline. A model’s relative AIC was quantified by subtracting the model’s AIC by RLWM’s AIC and dividing the answer by RLWM’s AIC. We also simulated data based on the models for validation.

#### 4.3.1 Two-parameters Reinforcement Learning (RL2)

The basic model was a reinforcement learning model (without the working memory component) with a delta rule learning. For each stimulus, *s*, and action, *a*, the expected reward was *Q*(*s*,*a*), and the Q value was updated with observing feedback, *r*_*t*_, through time. The Q values were updated based on a learning rate, *α*, and the difference between expected and observed reward at trial t (known as the prediction error: *δ*_*t*_ = *r*_*t*_ − *Q*_*t*_ (*s*, *a*)): *Q*_*t*+1_(*s*, *a*) = *Q*_*t*_ (*s*, *a*) + *α* × *δ*_*t*_. Choosing an action utilized the expected reward value. An action was probabilistically chosen, with a greater likelihood of selecting an action that had a higher Q value, using the SoftMax choice rule: *P*(*a*|*s*) = *e*^*βQ*(*s*,*a*)^⁄∑*_i_*(*e^Q^*^*β*(*s*,*a_i_*)^), where *β* is an inverse temperature free parameter. This model had two parameters of *α* and *β*.

#### 4.3.2 Four-parameters Reinforcement Learning (RL4)

This model in addition to RL2 includes a value for unrewarded correct responses and undirected noise in action selection. In this experiment, a correct response was sometimes rewarded and sometimes not rewarded. We estimated how much a person valued a correct response, irrespective of the reward by estimating the value for correct-but-not-rewarded items, i.e., *r*_0_. The model also considered an undirected noise, 0 < *ϵ* < 1, in the stochastic action selection, to allow for choosing an action that did not have the highest Q value. Accordingly, 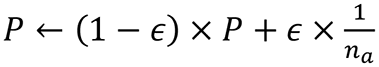 where 1/*n_a_* is a uniform probability of choosing an action.

#### 4.3.3 Reinforcement learning and working memory (RLWM)

We applied RLWM to estimate the working memory capacity and forgetting rate of the participants (Collins et al., 2017). This model had 8 parameters and consisted of two components. A working memory component with limited working memory *capacity, C*, and *forgetting rate,* ∅*_*WM*_*. The Q value was subject to decay with a forgetting rate, 0 < *ϕ* < 1, so for all the stimuli that are not current, *Q* ← *Q* + *ϕ* (*Q*_0_ − *Q*), where 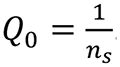.

The RL component had a *learning rate, α,* value for an unrewarded correct response, *r*_0_, undirected noise, *ϵ*, and a *forgetting rate, ϕ_RL_* (*β* was set constant at 100). We also allowed for the potential lack of an impact of negative feedback (*δ* < 0) by estimating a preservation parameter, *pers*. In that case, the learning rate is reduced by *α* ← (1 − *pers*) × *α*. Accordingly, *pers* near 1 indicated lack of an impact of negative feedback (learning rate close to 0; high preservation of Q value), and *pers* close to 0 indicated equal learning rate for positive and negative feedback.

The WM component was simulated as encoding of stimulus in a Q learning system, like the RL component but the outcome, *r*_*tt*_, was 1 for correct, 0 for incorrect (rather than the observed reward), the learning rate was set to 1 (*α* = 1), and at most *CC* stimuli could be remembered. We formulated the probability of a stimulus being in working memory as:

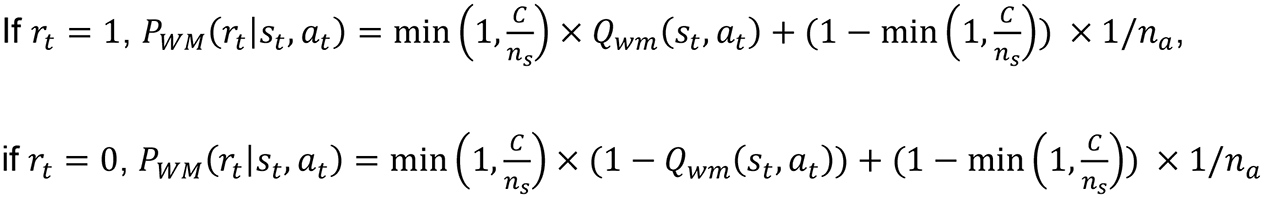

where *n*_*a*_ = 3 is the number of possible actions. In the RL case,

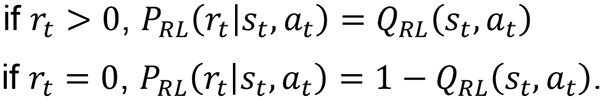

A *mixture weight*, *w*_0_, formulated how much each of the components was used for action selection. The weight was 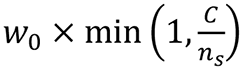 to represent the confidence in WM efficiency. This initialization reflects that participants are more likely to utilize WM when the stimulus set size is low. The overall policy was:

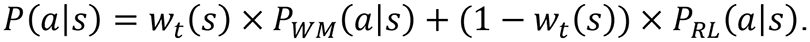

A Bayesian model averaging scheme inferred the relative reliability of WM compared with the RL system over time, *t*:

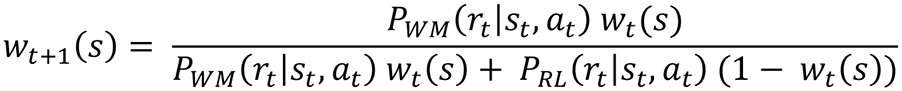

where *P_WM_* is the probability that action *a* is selected for stimulus *s* according to the WM component at time *t* and *P_RL_* is the probability of action selection according to the RL component. We assumed that although the *w*_0_ is the same for all stimuli, the development of mixture weight over time would be different for each stimulus because the probability of retaining a stimulus in working memory or another retention system is not equal.

### 4.4 Cross-task comparison

We determined the correlation between the logistic regression-estimated Beta values (from the trial-by-trial modeling of choices) for the three parameters - delay, number of previously correct answers, and set size - and their interactions and the number of events segmented. This analysis led to 6 correlations. We adjusted the significance level using Bonferroni correction to 0.0083. This analysis was replicated for the three movies.

We also evaluated a relationship between the number of events and working memory capacity and forgetting rate that were estimated by the RLWM model using a linear and quadratic model fitting. The models were compared using AIC. We report the difference between the linear and quadratic AICs – larger positive difference means that the quadratic model was a better fit, despite the increased number of parameters. We hypothesized that the number of events would reveal the estimated working memory span.

We ran an analysis on the subsequent memory performance of two extreme groups with fine (one standard deviation more than the mean) or coarse (one standard deviation less than the mean) segmentation to determine their different strategies. We refer to these participants as coarse- and fine-segmenters. Our hypothesis was that the coarse-segmenters and fine-segmenters used different long-term memory strategies to compensate for fast working memory forgetting rate. We examined the temporal order recognition performance for the linear movie to determine which group relied on the story structure for memory performance. We also split the temporal order tests into close and far temporally distant scenes to determine the effect of storyline. We predicted that temporal order recognition was due to the storyline if the effect was driven by the accuracy of remembering the order of far scenes. We then used natural language processing algorithms to count the number of “realis” events (i.e., factual and non-hypothetical words) and the total number of written words in the free recall task to determine which group recalled more events about the movies (Sap et al., 2020; Sims et al., 2019).

#### 4.4.1 Cross-task results without RLWM

In experiment 1, we studied a link between the effects of working memory on learning and event segmentation. The results without reinforcement learning modeling also showed a link between working memory limitation and event segmentation. Overall, participants learned the stimulus-action associations. For all set sizes, the accuracy of the last two iterations was on average more than 90% (M= 93.5%, SD = 3%; Figure S1A); however, the accuracy decreased with increasing set size (r = −0.94, p = 0.018). We analyzed the trial-by-trial performance with respect both to set size and to the maintenance of correct associations across intervening trials of a stimulus. The result of a multinomial logistic regression revealed that performance was reduced with increasing set size (t = −10.12, p < 0.001) and delay (the number of trials since a correct response to the current stimulus (t = −8, p < 0.001). By contrast, the performance improved with increasing total number of previously correct responses to the current stimulus (t = 5.95, p < 0.001). The interactions between set size and delay (t = −8.29, p < 0.001), set size and previous correct responses (t = 4.19, p < 0.001), and delay and previous correct responses (t = 5.18, p < 0.001) also affected the performance (Figure S1B), consistent with the previous report (Collins et al., 2017).

We observed a linear correlation between the total number of determined events and the beta-value for the interaction between set size and previously correct responses (ranked r = 0.34, p = 0.04). The result was similar for each movie separately (mixed: ranked r = 0.31, p = 0.064; non-linear: ranked r = 0.45, p = 0.006; linear: ranked r = 0.32, p = 0.057). This interaction indicates that learning is slower with higher set sizes, revealing a working memory limitation on association learning. However, the beta value does not clarify what aspect of working memory is correlated with the number of segmented events. Using the RLWM model we systematically tested for the relationship between working memory forgetting rate and capacity and the number of segmented events.

### 4.5 Figure

**Figure S1.**
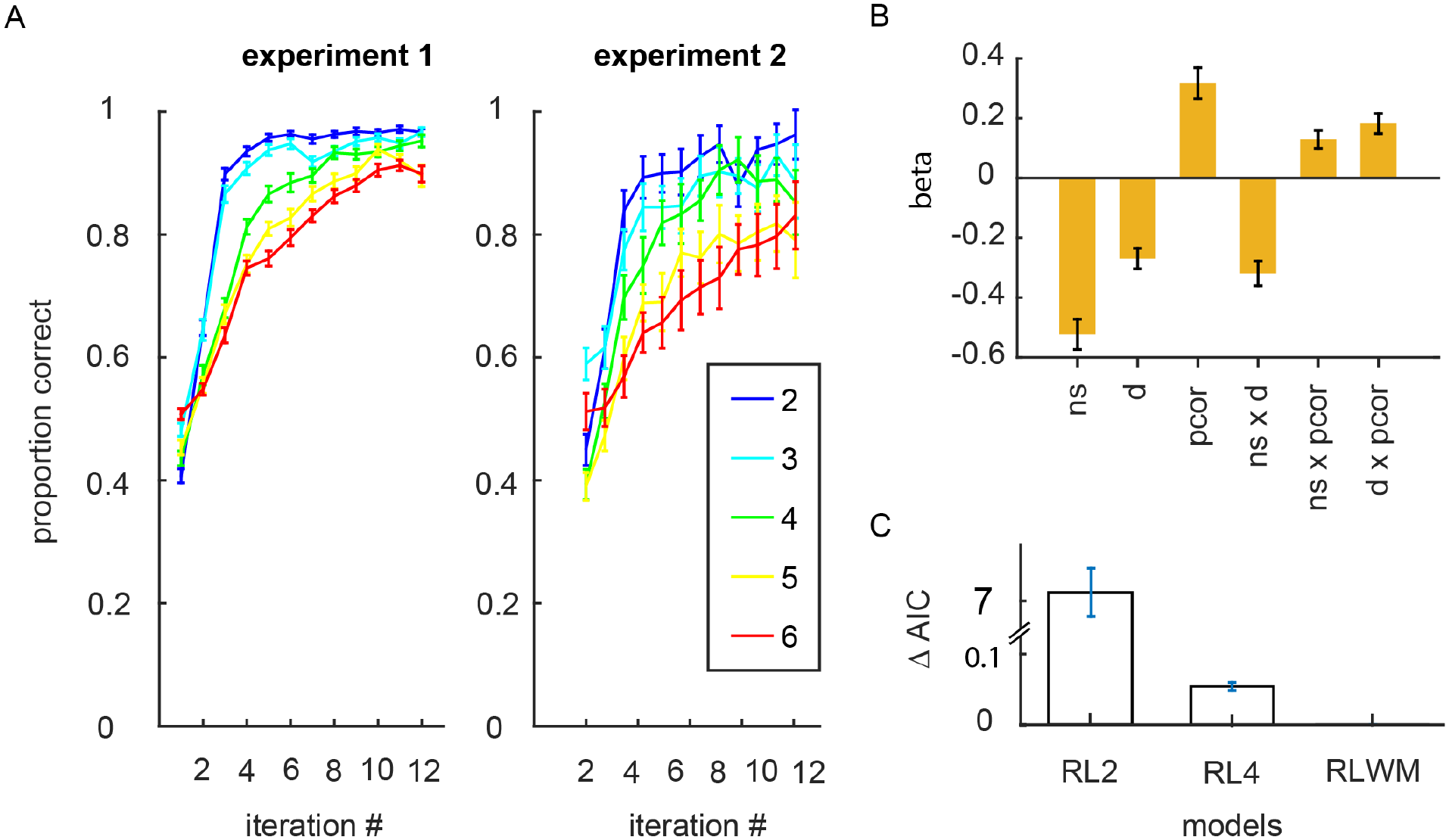
Association learning results: (A) Participants’ learning performance in experiment 1 (in left) and the learning performance in experiment 2 (in right). Learning was slower with increasing set size (ranging from 2 to 6). (B) The stimulus set size (ns), number of trials since the last correct answer (delay; d), number of previously correct responses (pcor), and their paired interactions affected the trial-by-trial performance in the learning task (shown for experiment 1). (C) The AIC of all models were measured relative to the RLWM (RL2: M = 7.4, SD = 1.68 and RL4: M = 0.06, SD = 0.03; note that smaller AIC reflects better fit). Pairwise t-test showed that RL4 was a better model than RL2 (t(34) = −25.99, p < 0.001), and RLWM provided the best fit (RL2: t(34) = 25.9, p < 0.001; RL4: (t(34) = 9.63, p = 0.03), shown for experiment 1. Error bars represent the standard error of the mean.

**Figure S2.**
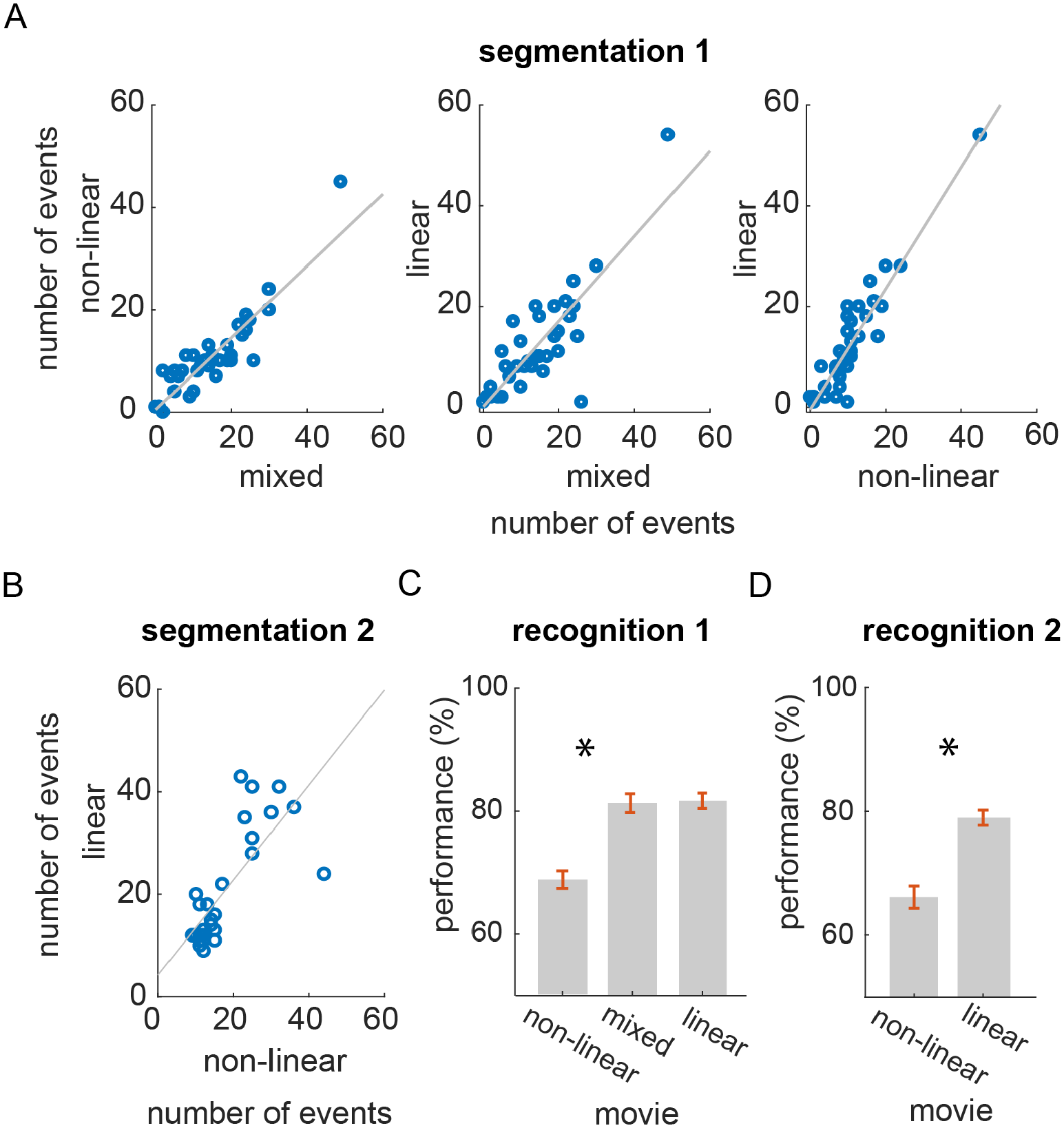
Event segmentation results: (A) Experiment 1: the number of events that participants segmented was constant across the movies. Correlation between the number of events for the mixed and non-linear movies (left), for linear and mixed movies (middle), and linear and non-linear movies (right). Each dot represents an individual participant; r values describe the ranked correlation coefficient. (B) Experiment 2: the number of determined events in the linear movie correlated with the non-linear movie. (C) Experiment 1: temporal order recognition performance was above chance for all three movies. The performance for movie non-linear movies was less accurate than for linear and mixed movies (* p < 0.05). (D) Experiment 2: Only subjects with above chance temporal order recognition were included in experiment 2. The performance for the linear movie was higher than the non-linear movie. Error bars represent the standard error of the mean.

### 4.6 Table

**Table S1.** Linear and quadratic model comparisons including all participants (except the one with outstanding number of events; n = 35). AIC difference represents the difference between the linear and quadratic AICs. * p-value < 0.05

| **Movie** | model | $R^2$ adjusted | p-value | AIC | AIC difference |
| --- | --- | --- | --- | --- | --- |
| All | Linear | -0.026 | 0.72 | -15.08 | 7.04 |
|  | Quadratic | 0.18 | **0.015 \*** | -22.12 |  |
| Mixed | Linear | -0.026 | 0.735 | -15.07 | 6.66 |
|  | Quadratic | 0.173 | **0.018 \*** | -21.74 |  |
| Non-linear | Linear | -0.02 | 0.581 | -15.27 | 7.14 |
|  | Quadratic | 0.189 | **0.013 \*** | -22.42 |  |
| Linear | Linear | 0.021 | 0.88 | -14.97 | 4.92 |
|  | Quadratic | 0.128 | **0.041 \*** | -19.90 |  |

**Table S2.**
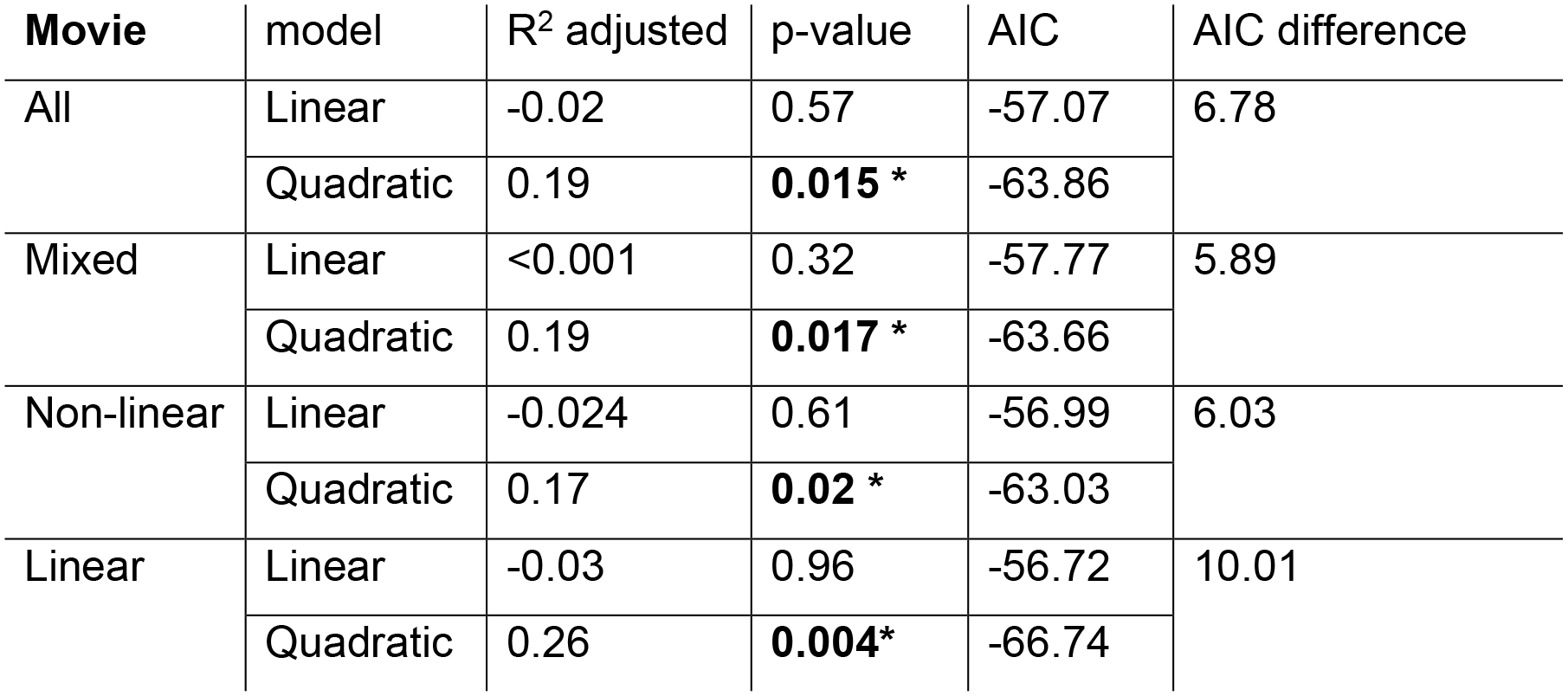
Linear and quadratic model comparison including 32 participants. Beside the one subject with outstanding number of events, three participants were excluded because they had a WM forgetting rate that was two standard deviations larger than the mean (*ϕ_WM_* > 0.6) and a problematic model fit with very high estimated learning rate (*α* > 0.9, two standard deviations larger than the mean) and low mixture weight (*w*_0_< 0.6), indicating that the working memory module of the model did not function in a regime representative of cognitive working memory function. * p-value < 0.05.

| **Movie** | model | R<sup>2</sup> adjusted | p-value | AIC | AIC difference |
| --- | --- | --- | --- | --- | --- |
| All | Linear | -0.02 | 0.57 | -57.07 | 6.78 |
|  | Quadratic | 0.19 | **0.015 \*** | -63.86 |  |
| Mixed | Linear | <0.001 | 0.32 | -57.77 | 5.89 |
|  | Quadratic | 0.19 | **0.017 \*** | -63.66 |  |
| Non-linear | Linear | -0.024 | 0.61 | -56.99 | 6.03 |
|  | Quadratic | 0.17 | **0.02 \*** | -63.03 |  |
| Linear | Linear | -0.03 | 0.96 | -56.72 | 10.01 |
|  | Quadratic | 0.26 | **0.004\*** | -66.74 |  |

**Table S3.** Linear and quadratic model comparison including the participant with the outstanding number of events (n = 33). * p-value < 0.05.

| **Movie** | model | R<sup>2</sup> adjusted | p-value | AIC | AIC difference |
| --- | --- | --- | --- | --- | --- |
| All | Linear | 0.23 | **0.002 \*** | -52.61 | 10.38 |
|  | Quadratic | 0.45 | **<0.001 \*** | -63.00 |  |
| Mixed | Linear | 0.22 | **0.003 \*** | -51.83 | 13.01 |
|  | Quadratic | 0.48 | **<0.001 \*** | -64.85 |  |
| Non-linear | Linear | 0.25 | **0.001 \*** | -53.40 | 8.95 |
|  | Quadratic | 0.44 | **<0.001 \*** | -62.35 |  |
| Linear | Linear | 0.18 | **0.007 \*** | -50.37 | 13.17 |
|  | Quadratic | 0.46 | **<0.001 \*** | -63.54 |  |

### 4.7 Movies stories

#### 4.7.1 Mixed

The animation depicted an overarching cliché love triangle story, along with some temporally interchangeable stories. It starts by showing a few animal couples going back and forth in a park. Then, there is a small lion that looks heartbroken. The lion sees a lioness, but there is a bigger lion that wants to meet the lioness too. The two lions fight for the lioness’s attention through a series of matches that have periods with an anticipated flow. For example, after each match, the score is shown on board. However, the score is not immediately shown after an eating contest. After the bigger lion wins the eating contest, it eats the small lion’s food too. Then the score is shown, and the matches continue. The small lion loses the competition and moves on to another part of the world. The end of the movie shows that the small lion meets a lioness.

#### 4.7.2 Non-linear

The animation is a sequence of independent events that involve crocodiles. It starts by showing two zebras listening to music next to a swamp. A crocodile suddenly eats one of them. Then, the crocodiles swim in a swamp just below the surface with only the eye visible. One of the crocodiles is wearing glasses. This crocodile stands up for a moment to clean its glasses and then it continues swimming below the surface. Next, a crocodile attacks a cow that is drinking water. The cow is too big for the crocodile so it cannot bite it. The cow, however, beats the crocodile in one attempt. Next, a crocodile is eating at a table in the swamp that has birds next to it. It puts catchup on the birds and eats them one by one by a fork. A bigger crocodile takes a cow into the swamp, but the cow defeats the crocodile and comes out. Next, a goat is swimming away from a crocodile, clearly scared. The swamp suddenly dries out. The crocodile cannot walk fast, but the goat happily leaves the swamp. Then, a crocodile attacks a cow that is by the swamp, but the cow skins the crocodile and takes it for tanning. Next, a crocodile with dental braces is shown drinking with a straw. A baby zebra plays by the swamp and bothers the crocodile. After that, a crocodile is shown participating in a non-violence resistance group with other animals, holding a peace sign. The movie ends with a scene of a very long crocodile on which a bird is happily picnicking.

#### 4.7.3 Linear

The movie depicts a linear life story of a pig. It starts by showing a caterpillar on a leaf. Then a big sow appears and gives birth to seven piglets. The piglets follow the sow in a line going around woods and crossing roads. Two of the pigs suddenly disappear; they were killed on the road. The rest of the piglets also disappear one by one, except for one. Then the caterpillar is netting a cocoon – showing the passage of time. The piglet grows up to be an ugly boar, and the sow is old. The sow dies. The pig meets three gilts. They reject him (depicted as a computer error message box) because he does not have money, he is ugly, and one of the gilts is already married. The cocoon is now complete, and the boar is still sad and alone. It bumps into a lion that was hunting for zebras. The lion gets happy for the catch, but the boar is too smelly. The lion puts the boar in a washing machine. It comes out as a red boar which is not desirable to the lion. The lion dumps the boar. The boar passes by the gilts again. This time, the one that was interested in a good-looking boar is interested and follows him.

### 4.8 Representative example of free recall

#### 4.8.1 Fine-segmenter

**Mixed**: giraffe dragging aligator along the ground. the painting was 2D but looked like the smaller tiger was flattened into it. smaller tiger takes a long time to eat. lots of couples at the beginning. bigger tiger cheats on the test. not sure what that map with the arrow meant but some cannon? but somehow smaller tiger crused bigger tiger yay

**Non-linear**: glasses aligator was funny. big bison/ox was hero of the story. wonder why that first zebra or animal just dove into the lake. aligator really wants to eat that bison. not sure what that wrench looking tool was, but aligator got skinned. part where the aligator puts sauce on the birds just standing there was unexpected but very funny

**Linear**: Time passing was kind of conveyed by the worm/caterpillar, which was really cool. lots of crisscrossing across the screen. what did the arrow signify? funny how the arrow lost the color in the wash and dyed him red. error boxes popping up was funny because it was almost like a computer game. not sure what happened to the other little babies along the way….

#### 4.8.2 Coarse-segmenter

**Mixed**: two different levels of lions competing with each other the lower level lion only won the writing test

**Non-linear**: aligators wants to eat other animals some animals sold the aligator skin

**Linear**: the washing machine switched the color of the pig and the color of the arrow the female pig fall in love with the male pig after he changed his color

## Notes

### Competing Interest Statement

The authors have declared no competing interest.

### Summary of Updates

We replicated the study in an independent participant sample. Considering the limitations imposed by the COVID-19 pandemic, we replicated the study remotely through an online setup. Although an online setup is not directly comparable to the well-controlled lab environment, we observed the key finding of the study in the online data. Namely, we observed the same direction of a link between working memory forgetting rate and event segmentation with the subsequent memory effect on temporal order memory. A caveat of the online study is that we could not identify coarse-segmenters in the data set acquired online. Instead of a U relationship, shaped by three groups of segmenters, we observed a correlation between two of the groups (mid and fine segmenters).

